# Simultaneous measurements of translation rate and transcriptome in single bacterial cells by scTR-seq uncovers linked regulation within a specialized active cell population

**DOI:** 10.1101/2025.04.22.649982

**Authors:** Christopher Baumann, Molly Hydorn, Samuel F. Cooke, Jonathan Dworkin, Adam Z. Rosenthal

**Affiliations:** Department of Microbiology and Immunology, University of North Carolina School of Medicine, Chapel Hill, NC; Department of Microbiology and Immunology, Vagelos College of Physicians and Surgeons, Columbia University, New York, NY 10032

**Author notes:** These authors contributed equally: Christopher Baumann, Molly Hydorn.

**Keywords:** translation rate, single cell RNAseq, phenotypic heterogeneity

## Abstract

Cell-to-cell variation within clonal bacterial populations provides bacterial communities important advantages including opportunities for bet-hedging and metabolic division of labor. In recent years the extent of bacterial heterogeneity has been documented both at the transcriptome level and with physiological measurements of cell growth rate and translation rate. However, methods that link physiological parameters to a single cell’s full transcriptomic state are lacking, making it difficult to identify the regulatory mechanisms that couple physiology and transcriptional output. Here we introduce a method that combines click-chemistry enabled labeling of nascent polypeptides to measure translation rates in single cells alongside microfluidic encapsulation and single cell transcriptomic measurements, providing a tandem measurement of translation rate and transcriptome in thousands of single *Bacillus subtilis* cells. In a culture experiencing nutrient limitation, we identified a subpopulation of cells with a higher rate of protein translation that uniquely overexpresses genes for several metabolic processes including acetoin production and arginine synthesis. Using a genetic approach informed by the gene expression in this subpopulation, we identified a regulatory mechanism that couples the increase in protein abundance of a transcriptional regulator AlsR with expression of *alsR* regulated genes in this subpopulation.

## Introduction

Morphological, physiological and transcriptional heterogeneity in bacterial cells exists even in clonal bacterial populations^1–5^ and provides several advantages to bacterial populations. For example, differentiation of a subset of cells into a specialized physiological state that is resistant to stresses, such as sporulation induced by starvation or temperature changes or antibiotic persistence, provides a bet-hedging strategy that maximizes population resilience by optimizing the growth strategy^6,7^. Additionally, differentiation of a clonal population into cell states that interact metabolically can provide a division of labor^8^. The metabolic interaction between differentiated sister cells can provide benefit to the community by bypassing metabolic incompatibilities^9^ and increasing biochemical reaction efficiency by optimizing resource allocation into different tasks within single cells^3^.

Many studies of bacterial cellular heterogeneity have relied on fluorescent transcriptional or activity reporters, which are typically limited to examining 4 or 5 spectrally distinct reporters at a time. Fortunately, recent advances in bacterial single-cell transcriptomics now enable the profiling of transcriptome-wide heterogeneity across thousands of cells^10–15^. While these new methods offer unbiased genome-wide insights, transcriptional variance may not always completely capture a cell’s physiological state. This is due to the low levels and short half-lives of mRNA in bacteria which, along with other factors, can result in inherently noisy and bursty transcriptional outputs. Thus, snapshots of the transcriptome may not fully represent the cell’s ongoing activity, but may instead reflect transient changes that do not drive sustained biological states requiring more sustained transcriptional coordination.

As single cell RNAseq measurements provide a snapshot in time^16,17^, inferring a physiological state with an inventory of transcripts does not always provide a straightforward interpretation. In both mammalian and bacterial systems, measurements of protein levels do not necessarily correlate to transcripts levels at the individual cell level^18,19^. To overcome these limitations, we sought to develop a tool that can combine single-cell transcriptomic data alongside single-cell measurements that report on other physiological activities within the same single cell, thus correlating the transcriptional inventory of each cell with a physiological signature.

In this study, we focus on coupling transcriptomic output with measurement of total protein synthesis rates (translation rates) because it is a fundamental measurement of bacterial activity. The recent development of techniques that assay single cell translation rates has revealed substantial cell heterogeneity in phylogenetically diverse organisms and under a range of growth conditions. For example, translation rates are bimodal in cultures of nutrient limited *Bacillus subtilis*^4^, as demonstrated by incorporation of a fluorescent tag into nascently synthesized proteins. This bimodality fits in well with other studies that find heterogeneity in translation^4,20,21^ and components that control translational output^5,22^. Here we set out to adapt the translational fluorescence measurement to instead generate a signal quantifiable by single-cell sequencing, so that it can be combined with transcriptomic signature in microfluidic single cell RNAseq barcoding approach we have recently described^10,23^. The simultaneous readout thus correlates transcription profiles with translation rate measurements in single cells and provides new insights about how cells link transcription and translational heterogeneity. We name this technique scTR-seq (**s**ingle **c**ell **T**ranslation and **R**NA **seq**uencing)

## Results

### Translation rate measurement using click-chemistry addition of oligonucleotide indexes

Methods to measure global translation rate are routinely used in mammalian cells and have been adapted for bacterial cells to measure overall translation rate^4,24^. These methods rely on the incorporation of a fluorophore proportional to the cellular translation activity. Several commercially available methods utilize incorporation of a click-chemistry enabled moiety into the elongating polypeptide either by including a click-enabled amino acid analogs (L-azidohomoalanine and homopropargylglycine, commonly abbreviated AHA and HPG^25^) or by adding O-propargyl-puromycin (OPP), a misincorporation signal containing an alkyne^26,27^. In bacterial systems several modified protocols of these click-based labeling methods are used for BONCAT (Bioorthogonal Noncanonical Amino Acid Tagging), a technique that allows labeling of specific proteins during synthesis using click-reagents^28^. We have previously utilized the OPP labeling technique to demonstrate that translation rate can vary amongst individual *B. subtilis* cells under certain growth conditions ^4^.

To develop a technique that links each cell’s translation rate measurement to other single cell sequencing outputs, we reasoned that substituting the incorporation of fluorophores with the rate-dependent incorporation of a user-defined oligonucleotide could provide a sequencing-based signal that can be captured via single cell sequencing. To achieve this, we adapted the OPP labeling method to incorporate an oligonucleotide probe (oligo^synth^) dual labeled with a click-chemistry-compatible azide molecule on the 3’ end and cyanine-3 (cy3) fluorophore on the 5’ end (IDT, see methods and Figure 1A).

**Figure 1:**
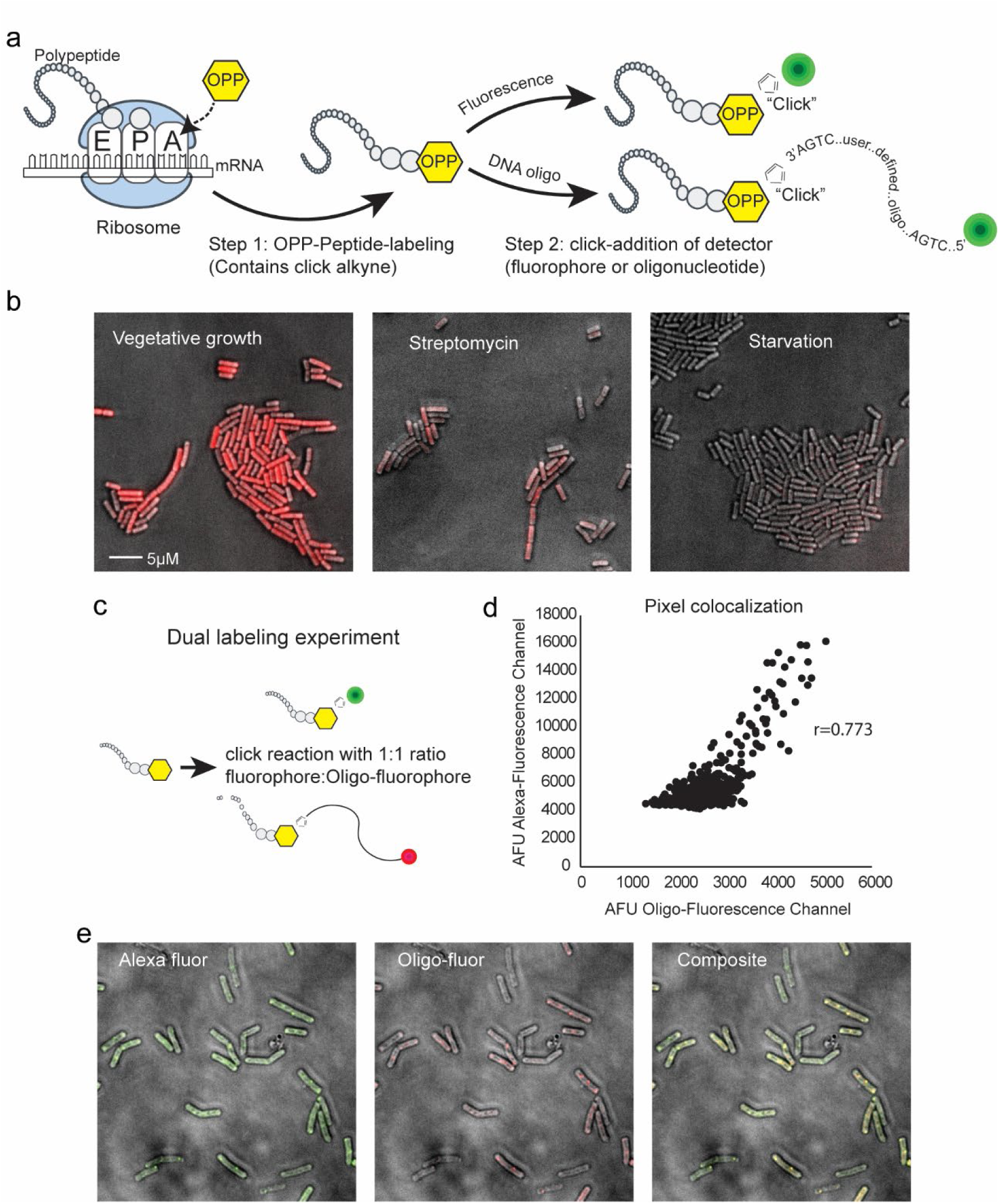
Click-chemistry based OPP labeling of protein synthesis using DNA oligonucleotide probes. **a.** Schematic representation of the OPP labeling method. **b.** representative images of *B. subtilis* cultures grown under different conditions and labeled with fluorescent and an azide-containing DNA oligonucleotide (oligo^synth^) using the OPP method. **c.** Schematic of the dual-labeling experiment. OPP was added to the culture using an equimolar ratio of fluorophore and oligo-fluorophore. **d.** Pixel colocalization from cells in the dual label experiment shows strong correlation between the fluorophore and the oligo-fluorophore signals. **e.** Representative images of the dual labeling experiment.

We cultivated *B. subtilis* cells in S7 media^29^ (see Methods) and at mid-exponential growth added a sub-inhibitory concentration of OPP for 20 minutes, as used in past bacterial work by us and others^4,24^. After cell fixation we attempted to incorporate oligo^synth^ using buffers provided in the commercial OPP-labeling kit (ThermoFisher), but our initial attempts did not provide noticeable labeling. We reasoned that this was due to the difference in azide groups in the kit-provided fluorophore and the azide-containing oligo^synth^, which may require different amounts of copper catalyst for efficient incorporation. To determine if copper addition could restore reaction efficiency, we designed a labeling buffer that includes BTTAA, PEG8000, copper sulfate and sodium ascorbate (see Methods). With this copper-containing buffer we were able to achieve observable labeling of cells with oligo^synth^. Using these optimized reaction conditions, we next compared the click-oligo labeling signal from cells grown in LB broth to mid-logarithmic growth (OD_600_= 0.7-0.8) to cells in the same condition that were exposed to translation inhibiting antibiotics (streptomycin, 15 minutes at 64ug/ml – a concentration used to classify streptomycin resistance^30,31^) or to cells undergoing acute starvation (cells resuspended in PBS). As expected, incorporation of oligo^synth^ was dependent on active protein synthesis (Figure 1B). As a control we compared oligo^synth^ labeling to traditional fluorophore signal by incubating OPP treated cells with both labels in our optimized copper-containing buffer (Figure 1C). This dual-label experiment confirms that the incorporation of the click-functionalized oligonucleotides oligo^synth^ measures translation rates in a manner similar to the traditional incorporation of fluorophores (r=0.773; Figure 1D and Figure 1E).

### scTR-seq: Pairing single cell translation rate and transcriptome sequencing using microfluidic encapsulation

To make oligo^synth^ compatible with our recently developed probe-based bacterial single cell RNA sequencing (ProBac-seq)^23^, we designed a probe complementary to our click-oligonucleotide incorporated using OPP-click addition. We included features on this probe that are compatible with the single cell indexing and library preparation workflow of ProBac-seq (Figure 2A and 2B). We then obtained combined single cell signals of both transcription and translation rate during the transition phase of the growth curve (300 minutes, Figure 2C) where fluorescence imaging of OPP-labeling indicated translation-rate heterogeneity. Cells were exposed to OPP for 20 minutes, fixed in formaldehyde and treated with the oligo^synth^ according to our optimized protocols. After click-incorporation, an aliquot of the cells was imaged, confirming single cell suspension and translation-rate heterogeneity, as reflected by different fluorescence intensity in individual cells (Figure 2D). The remaining cell sample was then incubated with transcriptome-wide probes (29,765 unique probes from the *B. subtilis* ProBac-seq probe-set^10^) and a probe complementary to oligo^synth^ and processed using ProBac-seq protocols^23^ with slight modifications (see Methods).

**Figure 2:**
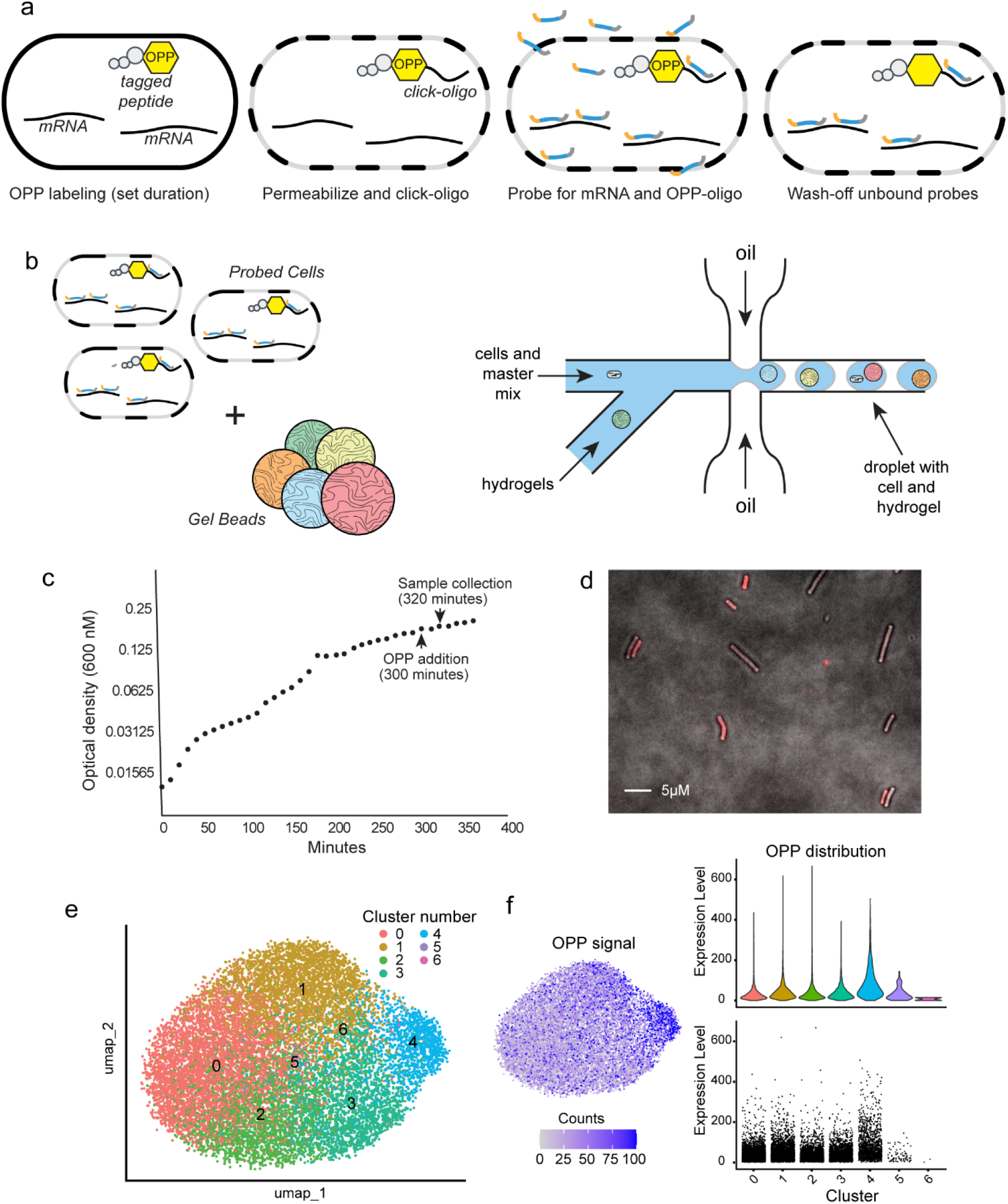
scTR-seq provides a tandem measurement of RNA and translation rate in single cells. **a.** Cells are incubated with OPP for a given duration of time, then fixed, permeabilized and conjugated with oligo^synth^ DNA oligonucleotide. OPP-labeled cells are then hybridized with probe sets complementary to the mRNA and OPP oligo sequences. Unbound probes are extensively washed to prepare labeled cells for single cell encapsulation. **b.** Cells and barcoded beads are separated into single cell emulsions using microfluidic single cell encapsulation. **c.** OPP was added to cells at 300 minutes, during the transition stage of growth, and a sample for analysis was harvested and fixed 20 minutes after OPP addition. **d.** A representative image from the sample after oligo^synth^ DNA oligonucleotide conjugation. **e.** UMAP two-dimensional representation of the cell clusters **f.** OPP translation rate projected on the UMAP. OPP distribution signal in each cluster is displayed in both violin plot (top) and corresponding cell plot (bottom).

Single cell encapsulation and sequencing resulted in 14,785 cells after filtering (Supplementary Table 1, methods). Clustering based on RNA expression patterns identified 7 unique cell clusters ranging in size from 5016 cells in cluster 0 to only 2 cells in cluster 6 (Figure 2E, and Supplementary Table 1). When overlaying the signal from the OPP probe with the RNA clustering, a significant enrichment of OPP signal in cluster 4 was noted (Figure 2F), indicating that this cell population had higher translational activity, consistent with the fluorescence observation of bright cells microscopically (Figure 2D).

Differential gene expression analysis identified several transcriptomic signatures that can distinguish cell clusters (Figure 3, Supplementary Figure 1). Cluster 4, which is associated with high translational activity, contains cells overexpressing genes involved in arginine biosynthesis, overflow metabolism (including acetoin production genes *alsS* and *alsD* and lactose lactate metabolism gene *ldh*), proton motive force amongst other processes (Figure 3a, Supplementary Table 2). In the context of *B. subtilis* physiology, our groups and others have previously identified several of these processes as heterogeneous in both transition growth phase^3,10^ and biofilms^32,33^. In addition to the enrichment of these functional genes, gene-set enrichment analysis (GSEA) also implicates a transcriptional repressor (Rex) and a transcriptional activator (AlsR) as potential genetic regulators controlling gene expression in this cell cluster (Figure 3). Two of the most differentially expressed genes in this cluster, *alsS* and *alsD*, encode the components needed for biosynthesis of acetoin, a key overflow metabolite produced by *B. subtilis* during the transition phase of growth and in biofilms. We have previously found these genes to be heterogeneous at the transition phase of growth using a different growth media^3^. Using them as a marker for the cell population in cluster 4 have used a fluorescent reporter for the promoter of the *alsSD* operon to confirm the heterogeneity of these genes in our culture conditions (Figure 3B).

**Figure 3:**
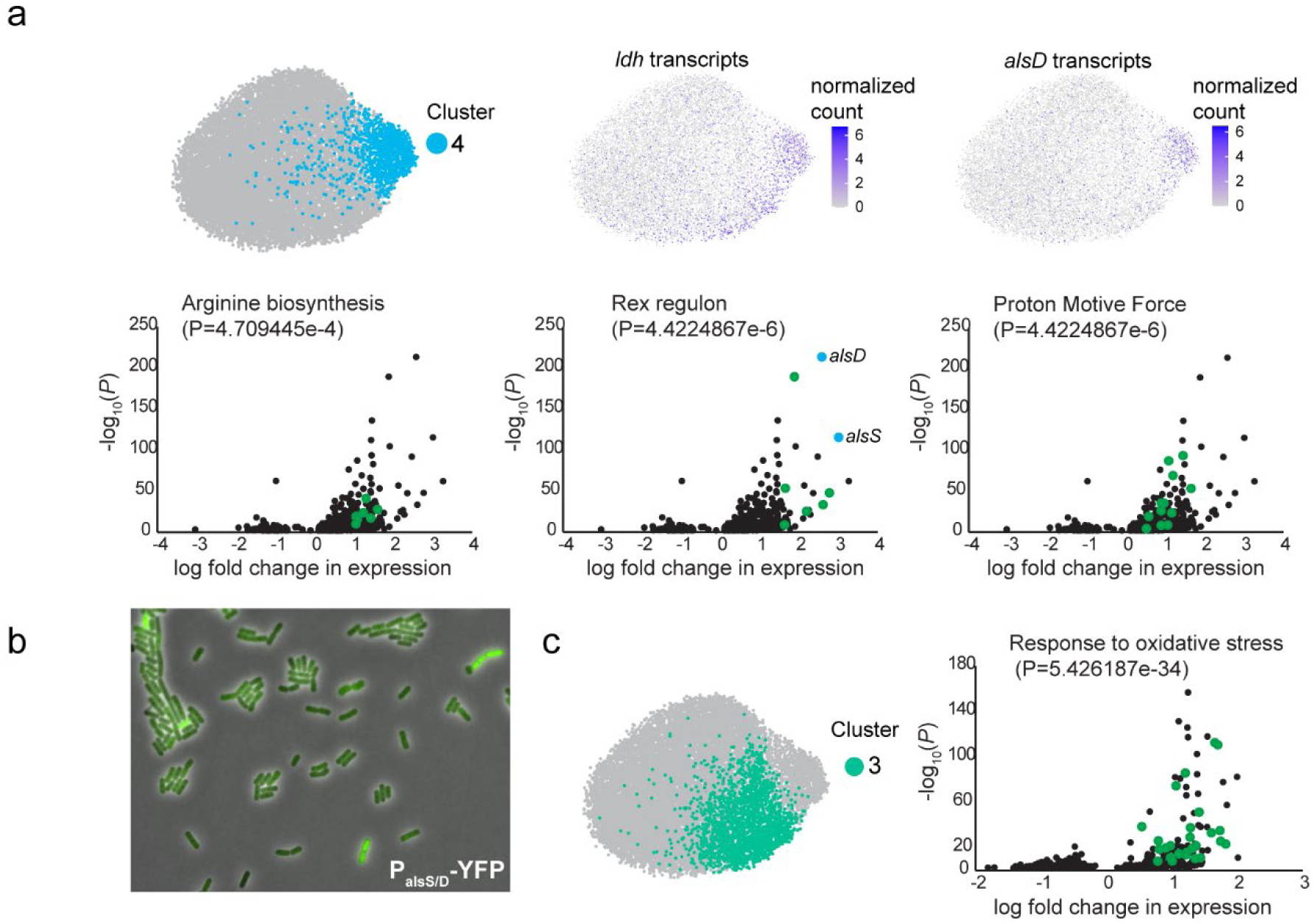
Distinct transcriptomic profiles and specific genetic regulators correlate with high-translation rate in single cells. **a.** Cells in cluster 4 (1,266 cells) and normalized single cell expression of overflow metabolism genes *alsD* and *ldh*, upregulated in cluster 4 (top). Volcano plots of gene sets enriched in cluster 4 including arginine biosynthesis, the Rex regulon and PMF genes (bottom). The *alsS* and *alsD* values in the Rex volcano plot are colored in blue because the *alsSD* promoter has an imperfect Rex consensus motif with low binding affinity. Overexpressed genes verified in these gene sets are colored green, *alsS* and *alsD* are putative Rex regulated genes and are colored in blue. **b.** A P*_alsSD_*-YFP reporter is heterogeneous in S7_50_ media. **c.** Cells in cluster 3 (teal, 2214 cells) are highly enriched in genes involved in the response to oxidative stress (green dots in the volcano plot). All adjusted *P* values are corrected by the method of Benjamini and Hochberg.

In addition to cluster 4, other cell clusters were also differentially enriched in distinctive functional gene sets. Specifically, Cluster 1 was enriched in amino acid synthesis genes, ribosomal subunits and other metabolic processes associated with cell growth and genes for motility and other cellular functions including purine synthesis (Supplementary Table 3). Cluster 3 was enriched with a large set of genes involved in response to oxidative stress (Figure 3C, Supplementary Table 4).

### Abundance of the transcriptional regulator AlsR in the high-protein-synthesis cell cluster correlates with high expression of AlsR target gene

To investigate the transcription of genes associated with the high translation activity of cells in cluster 4 specifically *alsS* and *alsD*, previously demonstrated to be heterogeneous ^3,32,33^, we examined the role of regulators implicated by GSEA. For example, *B. subtilis* Rex is a repressor that changes conformation in response to binding NADH and may regulate *alsS* expression^34^. To examine the role of Rex in our experimental context, we compared single cell transcriptional output from an *alsS* fluorescent promoter reporter (P_alsS_-YFP) in both wildtype *B. subtilis* and a strain in which Rex was deleted (Supplementary Table 5). The transcriptional output of *alsS* displayed cell-cell heterogeneity in both strains demonstrating that Rex is not required for formation of cluster 4 (Supplementary figure 2).

Heterogeneity in *alsS* expression could arise from heterogeneity in levels of AlsR, the transcriptional activator of the *alsSD* operon^35^. Specifically, cells enriched in acetoin biosynthesis genes also exhibit higher OPP probe signal (Figure 2F and Figure 3A), suggesting that differences in translation rates drive heterogeneity in AlsR protein levels and thereby differential expression of *alsS*. To assess this possibility and measure AlsR levels in single cells, we engineered a strain expressing an AlsR-GFP fusion at the native locus. This construct complements AlsR function since activity of P*_alsS_*-firefly luciferase transcriptional reporter was similar in AlsR and AlsR-GFP expressing strains (Supplemental figure 3). We incubated the AlsR-GFP strain with OPP and labeled with the Click-iT AlexaFluor 594 picolyl azide. Following imaging (Fig. 4A) and analysis, we denoted the top 25% OPP-labeled cells as the “HPS” (high protein synthesis) group (Fig. 4B). We obtained a random sample of the entire population (of equal number, n = 298), denoting this group as “RS” (random sampling) (Fig. 4B). Comparison of the two groups revealed that the HPS exhibited significantly higher AlsR-GFP fluorescence than the RS group (p < 0.0001 Mann-Whitney U test), indicating higher AlsR protein levels (Fig. 4B). We then determined the correlation between AlsR-GFP and OPP signals in single cells, finding a positive correlation (R^2^ = 0.5571) between the protein synthesis rates (OPP labeling) and GFP (AlsR-GFP) protein levels (Fig. 4C). Thus, differences in protein synthesis are likely sufficient to account for *alsS* transcription heterogeneity.

**Figure 4:**
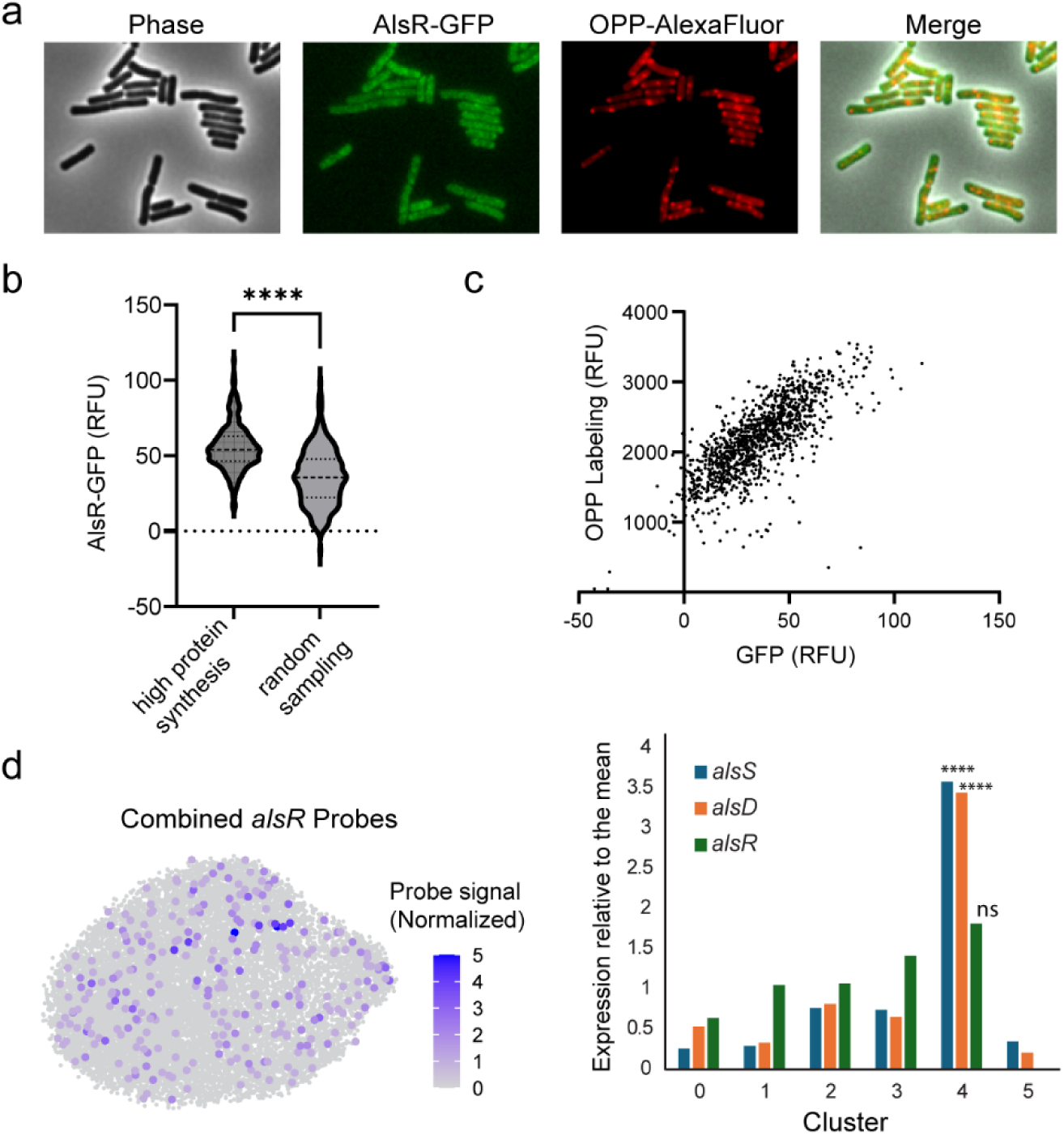
High protein synthesis rates account for increased AlsR protein levels. **a.** AlsR-GFP expressing cells (JDB4786) after OPP incorporation and AlexaFluor™ labeling shown in phase, GFP, mCherry, and merged channels. **b.** AlsR-GFP signal distribution in high protein synthesis cells (n = 298), defined by taking the top quartile of OPP-AlexaFluor fluorescing cells, and in a simple random sample (SRS, n = 298). ****, p < 0.0001 (Mann-Whitney U test). RFU = Relative Fluorescence Units **c.** AlsR-GFP signal (x-axis) and OPP-AlexaFluor signal (y-axis) for all cells in population (n = 1195). Linear regression analysis determined a line of best fit with parameters y = 20.02x + 1421, R^2^ = 0.5571. **d.** Combined single-cell probe signal from all *alsR* probes are highlighted on the UMAP. Cells with *alsR* signal were brought to the front and enlarged to aid visualization. ****Bonferroni corrected P value: *alsS* cluster 4 = 1.46E-11. *alsD* cluster 4 =1.37E-209. *alsR* signal is not statistically differentially abundant in cluster 4.

Although it is formally possible that *alsSD* operon expression heterogeneity is driven by differences in *alsR* transcription, the *alsR* gene does not exhibit statistically significant differential expression between clusters observed in scRNA-seq (Fig. 4D, Supplementary Table 6). This is consistent with data from an existing large gene expression study that reveal that the transcription of *alsS* and *alsR* across many conditions does not track: *alsS* transcript levels appear to be elevated in several conditions whereas *alsR* transcription remains essentially stable across conditions^36^ (summarized in supplementary figure 4).

## Discussion

Here we describe a method for simultaneously measuring bacterial transcriptomes and protein synthesis in single cells, thus connecting a physiological characteristic of a cell with its underlying gene expression. Our method (scTR-seq) adds to several multi-omic tools recently developed for single cell analysis including methods used in mammalian cells to associate transcription with chromatin accessibility^37^ and linking transcription with cell-surface-markers (epitopes)^38,39^. In most multi-omic methods the different omic signatures both represent static measurements, but the global protein synthesis measurement reported here quantifies the level of signal incorporated over a period of time, providing a biological rate that provides information about a cell’s global activity during a user determined timescale selected by the length of OPP incorporation. This timescale can be changed by altering the OPP “pulse” duration (incorporation time) to report on either instantaneous or long-lived measurements that average over a longer period. This user-defined decision should provide an indication of the time a given physiological state persists. This information is lacking in transcriptome measurements, especially in bacteria where mRNA half-life is on the order of minutes and transcriptional output is often bursty.

Our results identify a subpopulation of *B. subtilis* cells that have higher protein synthesis rates and differentially overexpress genes for several metabolic processes including overflow metabolism, amino acid biosynthesis and electron transport. We further demonstrate that the protein levels of AlsR, an important genetic regulator for key genes in this subpopulation, increases in this subpopulation even though it is not transcribed at significantly higher levels, offering a link between the protein-level abundance of a regulator and transcription of its downstream targets.

While the growth conditions assayed in this study identify highly active cells, the presence of individual bacterial cells with lower translation rates compared to most cells within a clonal population has been reported by several groups and is linked to antibiotic persistence^40–45^. Identifying and sequencing the transcriptome of these persistent cells could uncover regulators that lead to slow growth associated with antibiotic tolerance and persistence.

In addition to the phenomenon of persistence which links translational activity to antibiotic drug failure several classical studies have linked translational activity with cellular growth rates. This correlation was first observed in pioneering work from Ole Maaløe’s group describing a linear relationship between growth rate and protein synthesis per unit of RNA^46^. Using more sophisticated measuring methods and mathematical modeling later identified “rate laws” that describe this relationship between translational machinery and growth^47^. These mathematical relationships can correlate and rationalize global effects that couple the regulation of transcription to growth and translation^48^. However, some individual bacterial cells can have distinct growth rates^3,49–51^ and translation rates^4,5^ within the population raising the question of whether these cells obey the same growth laws, or whether states exist in which bacteria can decouple growth from protein production. If such cases exist, as we predict, a combined measurement of their transcriptome and translation rate will facilitate mechanistic investigation of such growth states.

This study uses click-chemistry incorporation of oligonucleotides to measure translation rate in single cells via sequencing. A recent study used click-based oligonucleotide attachment to measure nascent RNA in single eukaryotic cells using an adapted GRO-seq run-on transcription assay that labeled newly synthesized transcripts with a click-moiety^52^. In addition to these examples, common click-chemistry based EdU assay kits could be adapted to link DNA synthesis rates to single cell transciptomics and a click-based TUNEL staining kit can be used to measure the amount of DNA nicks and free 3’ ends. Together these physiological measurements can add context to transcriptomic signatures which can be hard to interpret due to dynamic factors including transcriptional burst kinetics.

## Supplementary Figure Legends

**Supplementary Figure 1: Heatmap of marker gene expression** (*z*-score of log-transformed values) organized into 7 clusters.

**Supplementary Figure 2: The transcriptional repressor Rex is not required for alsSD transcriptional heterogeneity.** An alsSD YFP promoter reporter signal is heterogeneous in both wildtype genetic background (left) and in a ΔRex background (right).

**Supplementary Figure 3: AlsR-GFP complements AlsR in regulating transcription of *alsS*.** Luminescence normalized by OD_600_ of P*_alsS_*-*firefly luciferase* (JDB4784, black) and *alsR*:: *alsR-GFP* P*_alsS_*-*firefly luciferase* (JDB4787, blue).

**Supplementary Figure 4: *alsR* transcript levels do not increase together with *alsS* transcripts in conditions where *alsS i*s highly activated.** A scatter plot showing the relative expression levels of either *alsR* (blue) or *alsS* (orange) obtained from normalized microarray dataset that includes data from 105 different conditions.

**Supplementary Figure 5: cell filtering cutoffs for single cell analysis.** Distribution plots of features (top) and unique molecular barcodes (bottom) used to determine cell cutoffs for single cell analysis. As demonstrated by lines on the right a cutoff was added to remove indexes with high signal that may be indicative of cell chains or clumps and a lower cutoff was applied to remove signal from noise that can occur from ambient probe signal in the samples.

## Supplementary Tables

**Supplementary Table 1:** Cell counts per cluster

**Supplementary Table 2:** Gene Set Enrichment Analysis for genes upregulated in cluster 4

**Supplementary Table 3:** Gene Set Enrichment Analysis for genes upregulated in cluster 1

**Supplementary Table 4:** Gene Set Enrichment Analysis for genes upregulated in cluster 3

**Supplementary Table 5:** List of bacterial strains used in this study

**Supplementary Table 6:** All differentially expressed genes in each cell cluster

**Supplementary Table 7:** Differentially upregulated genes in clusters 1, 3, 4 and 5 (for GSEA)

**Supplementary Table 8:** List of plasmids used in this study

**Supplementary Table 9:** Primers used in this study

## Acknowledgements

We acknowledge helpful advice from members of our laboratories. This work was supported by NIH R35GM141957 (JD) and startup funds from UNC Chapel Hill (AR).

## Materials and Methods

### Strain construction

Strains were derived from *B. subtilis* 168 *trpC2* except as noted and are listed in Table 1. Strains were constructed by transformation using conventional methodology and where necessary, media was supplemented with 100 µg/mL spectinomycin, 5 µg/mL chloramphenicol. Reporters were cloned into the *sacA* locus using pSac-cm^53^ and confirmed via whole plasmid sequencing via Plasmidsaurus, and *sacA* integration was confirmed by assaying growth on TSS glc and nongrowth on TSS suc plates. The AlsR-GFP protein fusion construct was created using Campbell integration via a pKL147-derived plasmid with a 450 bp homologous sequence to the 3’ end of the *alsR* gene. The Rex deletion strain (Δ*rex*::*kan*, BKK05970) originated from a large genome wide knockout collection^54^ and was ordered from the Bacillus Genetic Stock Center (https://bgsc.org/) and verified by sequencing. P*_alsS_*-YFP reporter, containing P*_alsS_* (encoding the region between −15 to −500 nucleotides upstream of the start codon) was previously reported^3^ and integrated into the Δ*rex* using standard *B. subtilis* protocols^55^.

### Media and Growth conditions

Bacteria were grown in S7 minimal media unless otherwise noted. S7 comprises 1X S7 salt solution diluted in reagent grade water (Teknova, W0225), supplemented with 1% glucose, 0.1% glutamic acid, 0.01% Casamino acids, 50mM NaCl, 40µg ml^-1^ tryptophan and 1X trace metals. The 10X S7 salt solution is prepared by dissolving 50mM MOPS, 10mM (NH_4_)_2_SO_4_, and 5mM KH_2_PO_4_ in ddH_2_0, buffered to pH 7.0 with 5M KOH, filter sterilized (0.22 micron) and stored at 4 °C. 100X trace metal solution contains 200mM MgCl_2_, 70mM CaCl_2_, 5mM MnCl_2_, 0.1mM ZnCl_2_, 100µg ml^-1^ thiamine HCl, 0.5mM FeCl_3_, & 2mM HCl (Teknova 2M2755). Cultures derived from a single colony were grown in S7 until OD_600_ = 0.3-0.4, then diluted to an initial OD_600_ read of 0.05 in S7. All cultures were grown in duplicate or triplicate.

### Luminescence

Luminescence assays were performed in a Tecan Infinite M200 Pro instrument with continuous shaking at 37°C, taking OD_600_ and luminescence reads every 10 minutes. Cultures were grown from single colonies grown overnight on LB plates at 37°C. Single colonies were picked into 2 mL S7 + CAA (1X MOPS (Teknova), 1.32 mM K_2_HPO_4_, 1% glucose, 0.1% glutamic acid, 0.01% casamino acids, 40 µg/mL l-Trp), unless otherwise stated, and grown at 37°C in a roller drum for ∼3.5 h. OD_600_ measurements were taken to ensure colonies remained in early exponential phase (between OD_600_ = 0.3-0.6). The cultures were then back diluted to OD_600_ = 0.05 in 0.5 mL S7 + CAA supplemented with 4.7 mM D-luciferin (Goldbio). Note that a spectrophotometer OD_600_ = 0.05 is ∼0.01 via plate reader. Cultures were aliquoted in triplicate in wells amounting 150 µL each in a 96-well white-walled, flat and clear bottom plate (Greiner Bio-One). Media only cells were used for background subtraction

### OPP labeling

OPP growth and labeling controls (Figure 1b and 1c) were prepared as previously reported^4^ with minor changes. Briefly, cells were grown to a mid-exponential OD_600_ = 0.7 before being separated into 450µL cultures, briefly subjected to an MIC_50_ concentration of streptomycin (64 mg/L) or nutrient-void media (1X PBS) or nothing (positive growth and click control) before addition of OPP and then cultured another 15 min in the OPP assay. One culture was left to grow without OPP for the 15 min to serve as a negative control for the click reaction. Cells were fixed by pelleting (6000xg for 5 min) and resuspension in 3.2% formaldehyde (final) in PBS for 10 min. Cells were then incubated in 0.5% TritonX-100 in PBS for 15 min for permeabilization. Cells prepared for scTR-seq cell-encapsulation (Figures 2 and 3) and the AlsR/OPP correlation experiment in figure 4 were grown on S7 minimal media from a culture re-diluted from an overnight culture originating from a single colony. 1000 µL cells at specified time points were transferred to disposable glass tubes. OPP was added to a final concentration of 10 µM. OPP incorporation was performed at 37 °C on a roller drum for 20 min and all subsequent steps were done at room temperature. Cells were fixed by adding formaldehyde to a final concentration of 1%. Cells were fixed for 30 min, harvested by centrifugation at 15,000 x *g* for 3 mins, and permeabilized using 100 µL of 0.5% Triton X-100 in PBS for 15 min. Cells were labelled using 100 µL of 1X Click-iT cocktail for 20 min in the dark. Cells were harvested and washed one time using Click-iT rinse buffer and then re-suspended in 50 µL of PBS for imaging.

### Oligo Click Reaction

Fixed and permeabilized OPP-treated cells were suspended in 200 µl of freshly made PEG-PBS click cocktail. This cocktail included 15% PEG8000 in PBS, 1 µl of the click-enabled oligo^synth^ oligonucleotide (5’ /5Cy3/TGT TCC CAA ATT GTA ACT TAT CGT TCC GTG AAG GCC AGA GTT ACT TCC CG/3AzideN/ 3’) (from a stock concentration of 100 µM, resulting in 0.5 µM final concentration), BTTAA (5 mM final) and CuSO4 (2 mM final) premix, and sodium ascorbate (100 mM final). Cells in the PEG-PBS cell suspension were left to incubate on a shaker at 50°C for 4 to 5 h. Cells were then washed twice in probe wash buffer before imaging or processing for scTR-seq.

### Click dual-labeling experiment

Dual-labeling reactions utilized the PEG-PBS click cocktail buffer used for Oligo-click reactions and included both the kit supplied fluorophore (Alexa-488) in the amount indicated in the commercial protocol (2.5 µl) and the Oligo^synth^ in the same concentration as for Oligo-click reactions (0.5 µm final).

### scTR-seq

Cells for scTR-seq were treated with OPP and click-coupled with oligo^synth^ as described in the Oligo-click reaction section. After oligo^synth^ addition cells were washed in PBS and a subset of cells was imaged to ensure cell integrity (a lack of cell lysis or clumping). The remaining cell sample was washed twice in probe washing solution at 50°C (5 × SSC, 30% formamide, 9 mM citric acid pH 6.0, 0.1% Tween 20 and 50 ug ml^-1^ heparin^56^) to remove unreacted oligo^synth^. Cells were then washed again in PBS and resuspended in 200 µl of probe binding buffer (5 × SSC, 30% formamide, 9 mM citric acid (pH 6.0), 0.1% Tween 20, 50 ug ml^−1^ heparin and 10% low molecular weight dextran sulfate^56^) and incubated at 50°C. Probes targeting the transcriptome were added as before^10^ and a probe (5’ CGG GAA GTA ACT CTG GCC TTC ACG GAA CGA TAA GTT ACA ATT TGG GAA CAG CTT TAA GGC CGG TCC TAG CAA 3’) was added to probe against the oligo^synth^ sequence.

Probes were left to hybridize overnight at 50°C with shaking. After hybridization cells were washed and counted on a Guava Easycyte HT cytometer as previously described^23^. A master mix was prepared for single cell encapsulation by mixing 19 µl of bioRad 4x ddPCR Multiplex Supermix, 3 µl “inDrop” primer (10 µm 5’ GAT GAC CCG GTC CAT ACA 3’), 3 µl “clickseq indrop” primer (10 µm 5’ CGG GAA GTA ACT CTG GCC 3’), 2 µl Reducing Agent B (10X Genomics), 4.8 µl ddH_2_O. A targeted addition of 16,000 cells in 43.2 µl PBS were added to this mix just before loading 70 µl of the mix onto a 10X Genomics “Chip-G” Microfluidic encapsulation device that was pre-loaded with 10X V3.1 beads as per 10X instructions and protocols for ProBac-seq^23^.

The encapsulation reaction was processed on a 10X Chromium iX device using the default run settings for 10X Chip-G devices. After the run completed the encapsulated cell emulsion was transferred to a PCR tube and cycled as previously reported for ProBac-seq^23^. After the barcoding PCR reaction the reaction was cleaned as previously reported^23^ and split into 2 separate reactions to independently amplify the transcript-binding probes (done using traditional ProBac-seq protocols) or the Feature-Barcode containing probe bound to the OPP-incorporated oligo^synth^. For amplification of the OPP signal 17 µl of the reaction was added to 20 µl of Q5 Hot Start 2x Mastermix (NEB), 1.5 µl of “Feature Bc PCR2 Fwd” Primer (10 µM 5’ GTC AGA TGT GTA TAA GAG ACA G 3’), 1.5 µl of “Clickseq to V3” primer (5’ GAT GAC CCG GTC CAT ACA CGG GAA GTA ACT CTG GCC 3’). The PCR mix was thermocycled using the following program: (1) 98°C for 30 seconds (2) 98°C for 10 seconds (3) 62°C for 20 seconds (4) 72°C for 20 seconds (5) repeat steps 2-4 for 15 additional cycles (6) 72°C for 2 minutes (7) 4°C Hold. After this PCR the OPP library was amplified a final time using the same cycling conditions as in the standard ProBac-seq protocol except that the forward PCR 3 primer was substituted for “10X feature Barcode PCR3 (PCR3 FWD)” primer (5’ AAT GAT ACG GCG ACC ACC GAG ATC TAC ACT CGT CGG CAG CGT CAG ATG TGT ATA AGA GAC AG 3’). Final libraries were validated as reported for ProBac-seq libraries and mixed in a 10:1 molar ratio (transcript:OPP) before sequencing on an illumina Nextseq 2000 using the following paired-end sequencing parameters: R1 28bp, R2 90bp, i7 index 8bp.

### Single cell bioinformatic analysis

Fastq files for both transcriptomic and feature barcode libraries were processed using the Cellranger (version 8.0.0) software using the cellranger count command. The reference probe genome was generated using cellranger mkref as reported previously^10^. After UMI correction and single cell expression matrices were generated, the raw data was loaded into R and processed using Seurat (version 5.2.1). The raw count data barcodes were filtered to only those with feature counts between 160-512 and UMI counts between 220-1000 to exclude potential multiplets and noise. The cutoffs chosen can be visualized in supplementary figure 5. Probes were collapsed to gene level data by taking the total counts across all barcodes for each probe and selecting the probe with the max count for each gene. The data was then normalized and processed using standard practices in the Seurat package. Data was log-transformed using NormalizeData (normalization.method = ‘LogNormalized’, scale.factor = 10000), and variable genes were selected using the FindVariableFeatures function. The highly-variable genes were z-scored with ScaleData. Linear dimensionality reduction using principal component analysis was done using RunPCA. UMAP projections were generated using RunUMAP(dims=1:30), nearest neighbors were determined via FindNeighbors(dims = 1:30), and Louvain clustering was performed using the FindClusters(resolution = 0.8) DGE analysis was performed using the FindAllMarkers function. Heatmaps were generated using the DoHeatMap function displaying the top 15 differentially expressed genes per cluster.

Geneset enrichment analysis was performed using curated gene lists and GO terms and P-value calculations accounted for Benjamini and Hochberg correction through the standard pipelines and annotations within the Bsub-cyc SmartTable gene set enrichment analysis function^57^.

### Microscopy and image analysis

Microscopy for figures 1-3 was done using agarose pads as previously described ^3^. Briefly, 1.5% agarose was prepared by adding 1.5% w/v low gelling agarose (LGA) into PBS. The solution was mixed and left at 65C overnight before use. From there, the 1.5% LGA in PBS remains at 65C. Pads were prepared by pipetting 1mL of the mixture onto a 22mmx22mm microscope cover slips and then putting another on top. The pads are left to solidify for at least 20 minutes. Pads were cut into desired sizes and 2 µl sample was pipetted onto each pad section. After 15 minutes of air drying, pads were inverted onto optical petri dishes (WillCo Wells B.V., The Netherlands). Dishes were imaged on an inverted Nikon Eclipse Ti2 microscope and imaged with a CFI PLAN FLUOR DLL 100X OIL objective, a D-LEDI light source and standard Nikon filter sets for GFP, RFP and far red fluorescence (Nikon catalog parts 96372, 96374 and 96376). Images were acquired using a Hamamatsu ORCA-Fusion BT Digital CMOS camera. Dual-fluorescence signal colocalization (Figure 1D) was calculated using the ImageJ tool EzColocalization^58^. Cells were segmented and overlapping pixel fluorescence intensity for each channel were calculated for calculation of the Pearson correlation coefficient.

Microscopy for figure 4 was performed using cells immobilized on 1% agarose pads prepared with PBS. Imaging was done with a Nikon 90i microscope with a Phase contrast objective (CFI Plan Apo Lambda DM ×100 Oil, NA 1.45), an X-Cite light source, a Hamamatsu Orca ER-AG, and the following filter cubes: GFP (ET Sputter 460/500×, Dm505, 510/10 m), and mCherry (ET Sputter Ex560/40 Dm585 Em630/75). The image stacks were analyzed in the software Fiji with the help of the MicrobeJ plugin^59^. The straighten and intensity options in the MicrobeJ plugin were used to measure the average fluorescence per pixel within each cell. Mann-Whitney statistical test was performed using MicrobeJ to assess significance of differences between the strains. A non-fluorescent control strain was used to subtract background and autofluorescence in each channel.

