## Supplementary figures and images for "Simultaneous measurements of translation rate and transcriptome in single bacterial cells by scTR-seq uncovers linked regulation within a specialized active cell population"

### Supplementary Figures

#### Supplemental 1

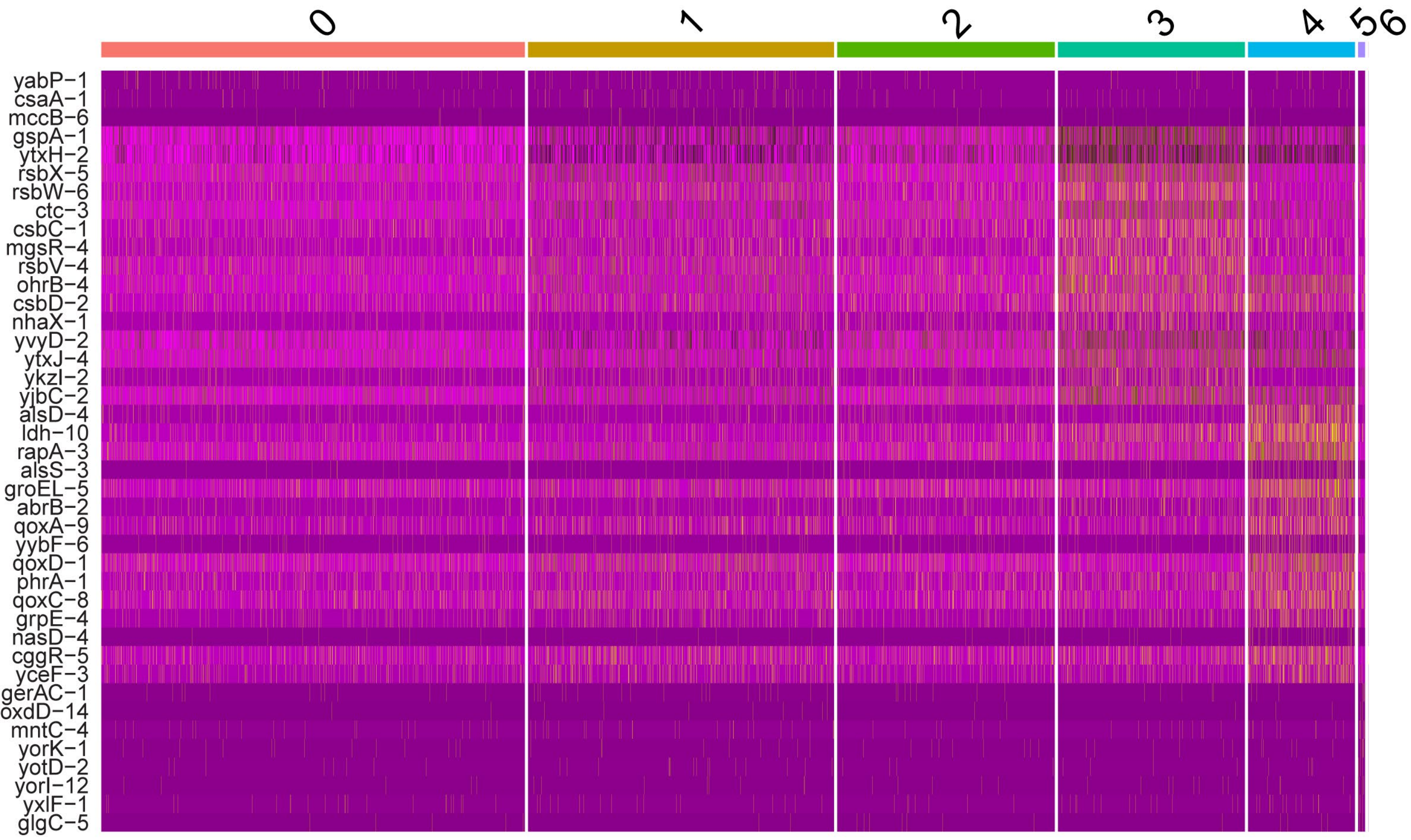

#### Supplemental 2

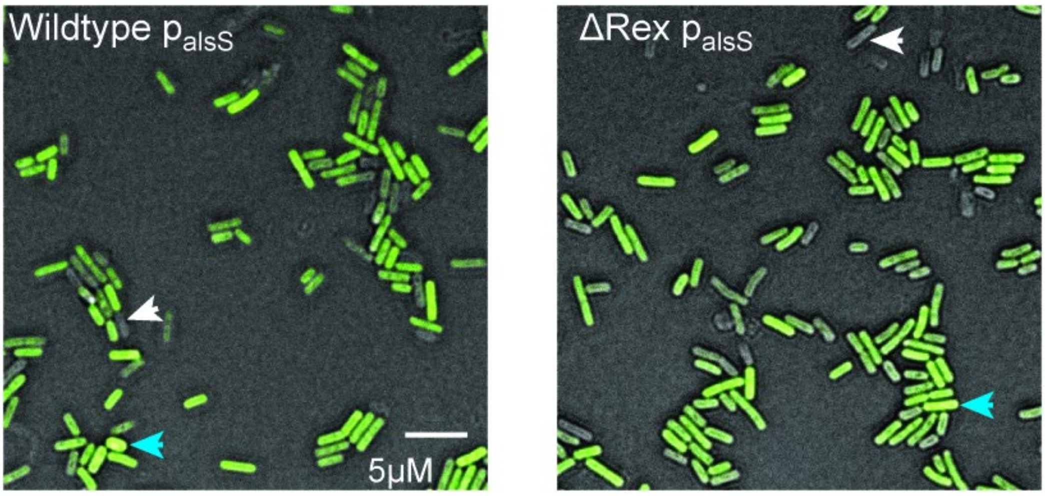

#### Supplemental 3

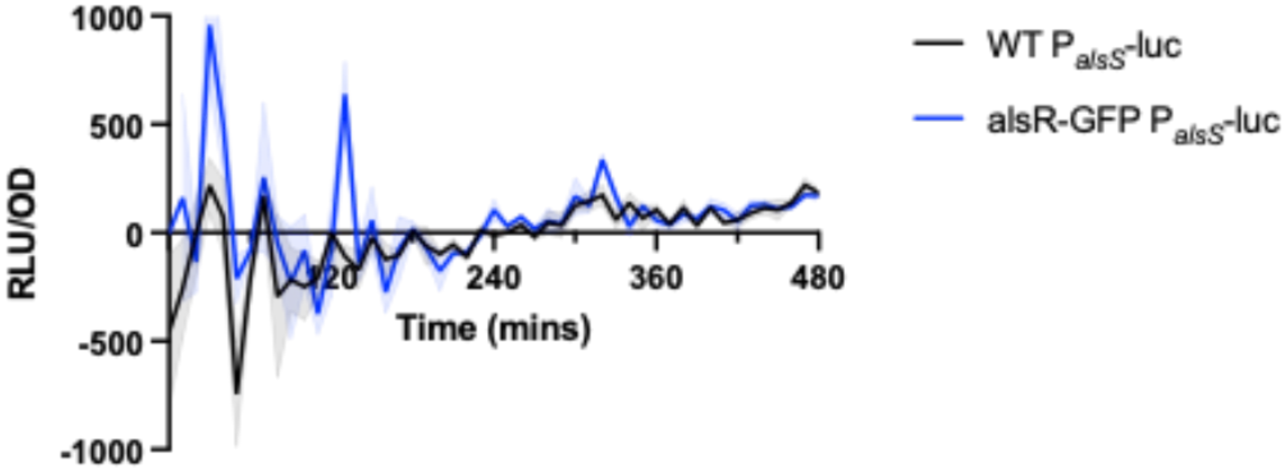

#### Supplemental 4

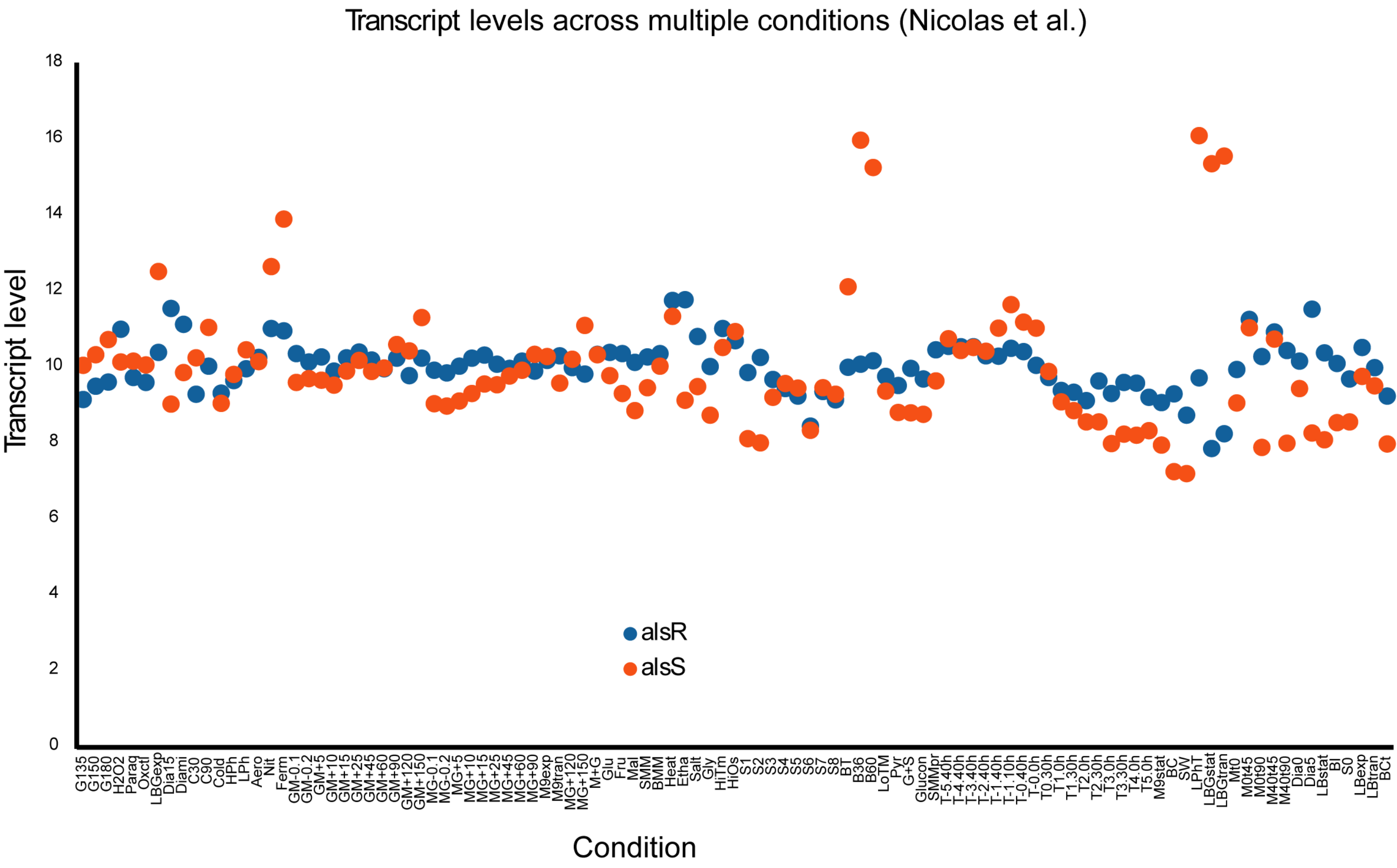

#### Supplemental 5

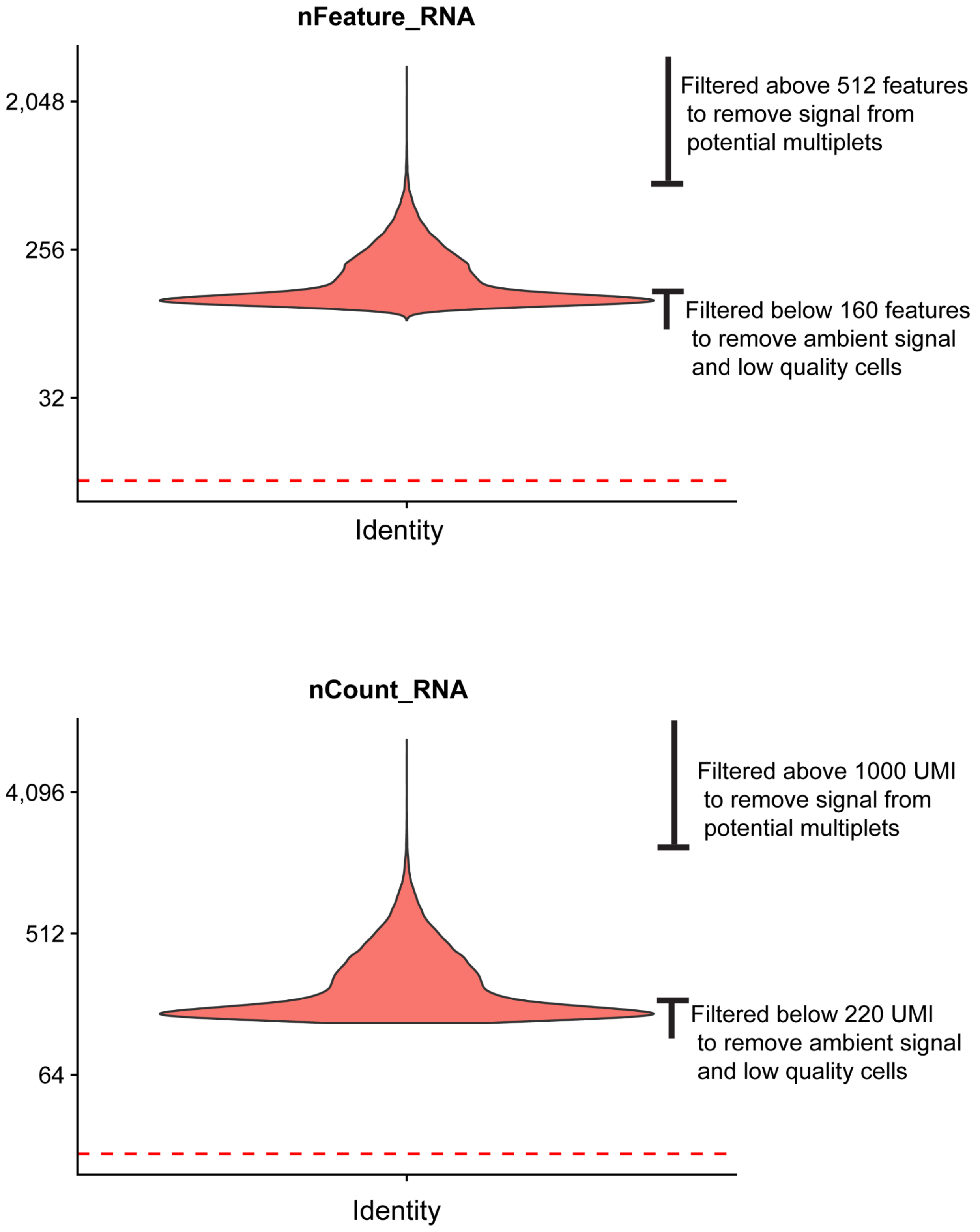
